# Combining organotypic tissue culture with multi-color fluorescence light-sheet microscopy (OTCxLSFM) – a novel tool to study glioma invasion/migration

**DOI:** 10.1101/2023.02.09.527810

**Authors:** Alicia Haydo, Andrej Wehle, Christel Herold-Mende, Donat Kögel, Francesco Pampaloni, Benedikt Linder

**Affiliations:** Experimental Neurosurgery, Dept. of Neurosurgery, Neuroscience Center, Goethe University Hospital, Goethe University Frankfurt, 60590 Frankfurt am Main, Germany; Division of Experimental Neurosurgery, Department of Neurosurgery, University Hospital Heidelberg, INF400, 69120 Heidelberg, Germany; German Cancer Consortium DKTK Partner site Frankfurt/Main, Frankfurt am Main, Germany and German Cancer Research Center DKFZ, 69120 Heidelberg, Germany; Buchmann Institute for Molecular Life Sciences (BMLS), Goethe University Frankfurt, 60438, Frankfurt am Main, Germany

**Author notes:** Correspondence: Benedikt Linder; 069 6301 6940.

**Keywords:** Glioblastoma, Light-sheet microscopy, Migration, Invasion

## Abstract

Glioblastoma is a very aggressive tumor and represents the most common primary brain malignancy. Key characteristics include its high resistance against conventional treatments, such as radio- and chemotherapy and its diffuse tissue infiltration, preventing complete surgical resection. The analysis of migration and invasion processes in a physiological microenvironment allows for enhanced understanding of these processes and can lead to improved therapeutic approaches. Here, we combine two state-of-the-art techniques, adult organotypic brain tissue slice culture (OTC) and light sheet fluorescence microscopy (LSFM) of cleared tissues in a combined method termed OTCxLSFM. Using this methodology, we can show that glioblastoma tissue infiltration can be effectively blocked through treatment with arsenic trioxide, as well as genetic depletion of the tetraspanin, transmembrane receptor CD9. With our analysis-pipeline we gain single-cell level, three-dimensional information, as well as insights into the morphological appearance of the tumor cells.

## Introduction

Tumors arising in the central nervous system are particularly devastating considering that the brain is one of the most delicate organs with a very special physiology including the blood brain barrier, its specialized cell types and its central importance preventing aggressive surgery. The most common and aggressive primary brain tumors are Glioblastomas (GBM), which are graded as a grade IV tumor according to the current WHO classification system (Louis *et al*. 2021). Key characteristics of GBM are their particularly high resistance against conventional chemotherapeutics and radiotherapy and their fast and highly infiltrative growth into the surrounding brain tissue, which frequently leads to tumor dispersal across the corpus callosum (Becker and Yu 2012), further complicating surgical removal and localized therapies such as radiation treatment. Despite an aggressive therapy consisting of maximally possible surgical resection followed by combined radio-chemotherapy with Temozolomide (TMZ), the median survival barely exceeds one year and the 5-year-survival rate is below 5% (Stupp *et al*. 2005; Stupp *et al*. 2007; Stupp *et al*. 2009; Luwor *et al*. 2013). Although novel treatment approaches such as Tumor Treating Fields show some success (Toms *et al*. 2019; Rominiyi *et al*. 2020), recurrences are common and overall survival is still particularly low. In fact, patients suffering from GBM have one of the worst prognoses compared to other primary cancers (Quaresma *et al*. 2015).

One major reason for failure of glioma treatment is the infiltration of tumor cells into the surrounding brain parenchyma. This inevitably excludes a total resection of the tumor and recurrence is virtually unavoidable due to the presence of treatment-resistant glioma stem-like cells (GSCs). It is known for many years that glioma cells migrate along existing anatomic brain structures like white-matter tracts, blood vessels and the extracellular matrix via interaction of multiple transmembraneous receptors (Ariza *et al*. 1995; Wiranowska *et al*. 2006; Krusche *et al*. 2016). Recently it has been reported that in particular white matter tract invasion of GSCs is regulated by a Notch1>Sox9>Sox2 feedback loop (Wang *et al*. 2019). Compared to the majority of tumor cells, these GSCs have, among other features, a higher differentiation potential, allowing them to replenish a tumor, usually reside within specialized niches (Gilbertson and Rich 2007; Soeda *et al*. 2009; Hide *et al*. 2019), and express various marker proteins associated with stemness (e.g. OLIG2, SOX2, SOX9) while lacking marker proteins of terminal differentiation like NeuN or MAP2 GSCs (Boumahdi *et al*. 2014; Trepant *et al*. 2015; Yoshimura *et al*. 2015; Bradshaw *et al*. 2016; Garros-Regulez *et al*. 2016; Tirosh *et al*. 2016; Voronkova *et al*. 2017).

Current approaches to study migration and invasion of GBM cells are either restricted to “classical” cell cultures of cells grown in serum-supplemented medium and consist of highly artificial scratch (“wound healing”) assays or invasion assays such as boyden chamber assay (de Gooijer *et al*. 2018). For once, these assays might have value in a very basic question, but they fail to provide the three-dimensional complexity and composition of a brain parenchyma. As such, even a boyden chamber assay, which employs Matrigel as a surrogate for an extracellular matrix (ECM), is more suited for epithelial cancers considering the composition of the ECM in the brain compared to epithelia. On the other hand allow complex model systems such as in vivo transplantation studies the most physiological model, but the study of invasion is usually restricted to staining of sequential sections and therefore very cumbersome, time-consuming and error-prone given that the sections need to be perfectly aligned. A rarely used, but powerful method consists of placing fluorescently labeled tumor cells onto murine brain slices and monitor tumor development for several days to weeks. We have successfully applied these so-called organotypic tissue slice cultures (OTCs) in previous studies and could show that this method has potential for further development (Remy *et al*. 2018; Linder *et al*. 2019; Gerstmeier *et al*. 2021; Linder *et al*. 2022). So far, we have analyzed tumor growth using an epifluorescence microscope and measured the tumor area on top of the brain slices. However, since GBM are known for their infiltrative phenotype and our current method lacks an efficient way to study invasion we combined the OTC-model with state-of-the-art multicolor fluorescence light sheet microscopy (LSFM) (Hotte *et al*. 2019).

Light sheet fluorescence microscopy (LSFM) is a three-dimensional imaging technique particularly suited for the analysis of large specimens, such as tumor biopsies, in their entirety (so-called *in toto* imaging) (Pampaloni *et al*. 2013). In LSFM, optical sectioning is achieved by illuminating individual planes of a specimen with a laser light sheet. The fluorescence emitted at each plane is collected by an objective lens placed perpendicularly to the optical axis of the light sheet. By translating the specimen through the light sheet, a three-dimensional data set is recorded. At variance with confocal microscopy, in LSFM only the plane which is imaged is excited by the light sheet. Thus, photo-bleaching and photo-toxicity in the specimen are reduced by several orders of magnitude compared to confocal microscopy. Moreover, the acquisition time is much faster compared to a standard confocal microscope (Stelzer *et al*. 2021).

The combination of LSFM and optical clearing is particularly powerful for digital pathology (Pampaloni *et al*. 2015). Optical clearing is obtained by homogenizing the refractive index in the specimen by using organic or aqueous solvents as index-matching media. With optical clearing, whole organs (including the human brain) become highly transparent. Therefore, even deep tissue regions are accessible to investigation with light microscopy (Hof *et al*. 2021). In general, optical clearing only works with chemically fixed samples. Current efforts aim to increase the throughput of the specimens that can be analyzed with LSFM and optical clearing, and to simplify specimen preparation. For instance, Glaser et al. 2022 developed an open-top LSFM for the fast imaging of very large, cleared organs (Glaser *et al*. 2022). We recently introduced thermoformed FEP-foil cuvettes for the multi-scale microscopy analysis of live and optically cleared large tissue specimens (Hof *et al*. 2021). The FEP-foil cuvettes allow for a straightforward and fast specimen preparation, which increases and improves the throughput of LSFM.

Here, we provide first data from our newly developed method OTCxLSFM, by combining a physiological tumor growth assay using OTCs with a microscope technique (LSFM) that is suited to measure large samples with high resolution after tissue clearing using CUBIC2 (Richardson and Lichtman 2015). As a model substance we employ arsenic trioxide (As_2_O_3_, ATO), which we recently employed to inhibit stemness in GSCs, when combined with the natural anticancer agent (-)-gossypol (Gos, also known as AT-101) (Voss *et al*. 2010; Meyer *et al*. 2018; Linder *et al*. 2019). We further provide evidence that this model system is also suited for genetically modified tumor cells by investigating the role of CD9 for migration and invasion.

## Materials and Methods

### Cell and Cell Culture

Human Glioblastoma stem like cells GS-5 GFP/Luc have been described previously (Linder *et al*. 2019) and are derived from GS-5 (Gunther *et al*. 2008), which were kindly provided by Katrin Lamszus (UKE Hamburg, Germany). NCH421k and NCH644 (Campos *et al*. 2010) were kindly provided by Christel Herold-Mende (University Hospital Heidelberg, Germany). The cells were cultured in Neurobasal medium (Gibco, Darmstadt, Germany). The medium was supplemented with 1x B27, 100 U/ml Penicillin 100 µg/mL Streptomycin (Gibco), 1x GlutaMAX (Gibco), 20 ng/mL epidermal growth factor (EGF, Peprotech, Hamburg, Germany) 20 ng/ml and fibroblast growth factor (FGF, Peprotech). HEK293T (ATCC #CRL-3216) were cultured in Dulbecco’s modified Eagle’s medium (DMEM GlutaMAX) supplied with heat-inactivated 10% FBS and 100 U/ml Penicillin 100 µg/ml Streptomycin (all from Gibco). Arsenic trioxide (As_2_O_3_, ATO; Sigma-Aldrich) was solved in 1 M NaOH, diluted with PBS (Gibco) to 0.5 M and solved at 80°C while stirring. The solution was then sterile filtrated and kept for long-term storage. An intermediate dilution of 15 mM was used for all consecutive dilutions.

### Synthesis of lentiviral supernatant and transduction

HEK293T cells were used to generate lentiviral particles. Therefore the cells were seeded in 6-well plates at a cell density of 300,000 cells and incubated overnight at 37 °C. Following, medium of the HEK293T cells was changed to transfection medium containing DMEM and 10% FCS (Gibco), incubating the cells for 30 min at 37 °C. Afterwards, transfection with the plasmid DNA was performed. For this, the DNA mix (1.34 μg shRNA-plasmid DNA (TRCN0000296954; TRCN0000291711, Sigma-Aldrich, Taufkirchen, Germany) or pLV[Exp]-EGFP:T2A:Bsd-CMV>ORF_Stuffer (VectorBuilder, Neu-Isenburg, Germany), 0.67 μg gag/pol plasmid (psPAX2, addgene #12260) and 0.86 µg VSV-G envelope plasmid (pMD2.G, addgene #12259) was prepared in 110 μl Opti-MEM, vortexed and incubated for 10 min. In this time the Fugene HD mix (8.6 μl Fugene HD (Promega, Madison, WI, USA) in 110 μl Opti-MEM was prepared and mixed thoroughly. After incubation, 110 μl of the Fugene HD mix was pipetted onto the DNA mix and incubated for 30 min at room temperature. Following the incubation, 220 μl of the mixture was added to the HEK293T cells. The plate was incubated for 6 to 7 hours at 37 °C. Afterwards, the medium was aspirated and refreshed using NB-A medium, following incubation overnight. After 16 h and 40 h the supernatants were collected, and pooled with the supernatants from the day before and afterwards filtered using a 0.45 μm filter and stored for a short time period at -21 °C or were used directly. Following, GSCs were used to generate stable shCD9 cell lines. As well as, after confirming efficient depletion of CD9 via western blot, generating GFP positive cell lines from the corresponding shCD9 cells. For this purpose, the corresponding virus was produced. Prior seeding the cells in a 24 well plate with a cell density of 500 000 cells per well, wells were treated with protamine sulfate to allow better adhesion onto the plate. Afterwards, the viral supernatants were added in a 1:1 dilution onto the corresponding cells. The cells were incubated for 72 h at 37 °C. After incubation, the media was changed to medium containing half the concentration of the selection antibiotic puromycin (Santa Cruz Biotechnology, Heidelberg, Germany)/ blasticidin (Thermo Fisher Scientific, Waltham, Massachusetts, USA). After 24 h of incubation the full concentration of the selection antibiotic (puromycin 0.5-2 μg/ml; blasticidin 1.25-2.5 μg/ml) was added at the cell line-dependent concentration. NCH421k and NCH644 were transduced with the virus made with the shCD9-constructs TRCN0000296954 and TRCN0000291711, respectively. The viability of the cells was checked every day for 4 days, containing each day a washing step. After 4 washing steps the cells were able to be transferred into S1 laboratories.

### Quantitative real time polymerase chain reaction

To analyze differential gene expression in GSCs, quantitative real time polymerase chain reaction (qRT-PCR) was performed. For this purpose, 300,000 cells were seeded in a 6-well plate in triplicates and were incubated for 48 h. Afterwards, cells were pelleted by centrifugation and washed with PBS. After this, the RNA was isolated directly. The isolation was performed according to the manufacturer’s protocol from the ExtractMe Total RNA Kit (7Bioscience, Hartheim, Germany). After isolation, the RNA concentration was measured using the Tecan Spark plate reader. The RNA samples were used directly for cDNA synthesis, using 1 µg of the sample. After this, the corresponding amount of DEPC-H2O was added. Adding 1 μl of 50 μM Oligo(dT)20, 1 μl Random Primer 150 ng absolute (3 μg/μl) and 1 μl 10 mM dNTP Mix to each of the RNA mixtures. The total volume was now set up to 14 μl. After pipetting, the mixture was transferred into an Eppendorf cycler (Eppendorf AG, Hamburg, Germany) and incubated for 5 min at 65 °C, followed by being cooled down on ice for 1 min. Followed by adding a mixture of the enzyme Superscript III (200 U; Invitrogen, Carlsbad, USA), DEPC-H2O, 0.1 M DTT and 5x first strand buffer. Thereafter, samples were heated up again for 5 min at 25 °C followed by a 1 h incubation at 50 °C and lastly a 15 min incubation at 70 °C. The resulting cDNA was diluted with 180 μl of DEPC-H2O and used directly for qRT-PCR. For each target gene the corresponding primer was added. To each master mix of the gene of interest, FastStart Master Mix and DEPC-H2O was added. This mixture was pipetted on a Micro-AMP fast 96-well reaction plate, as well as the desired cDNA. Afterwards, the plate was sealed with an adhesive film and was analyzed using the StepOnePlus qPCR machine (Applied Biosciences, Foster City, California, USA). Gene expression quantification using the ΔΔCT-method was performed using Graphpad Prism 9 (GraphPad Software, Inc., La Jolla, California, USA). Therefore, the samples were baseline-corrected by subtracting the housekeeper gene (TBP). Following with another baseline-correction by normalizing the samples towards the shCtrl cells. The fold change of each gene was generated by performing the equation: Y=2^-Y. The generated data was visualized, and ordinary one-way-ANOVA was performed to provide statistical information about the gene expression of each condition.

### Sphere Formation Assay

To determine changes in the sphere area/number and stemness character of GSCs after CD9-depletion, cells were seeded at a density of 500 cells per well in a 96-well plate. Each condition was seeded in 5 to 10 replicates. Following, the cells were incubated in an incubator at 37 °C for 7 d for the NCH cell lines. Pictures of the spheres were obtained with a Tecan Spark plate reader (Tecan, Männedorf, Austria), which was equipped with a camera. After this, the pictures were evaluated using Fiji (Schindelin *et al*. 2012). The spheres were counted with a self-made macro for Fiji. The mean sphere area as well as the number of spheres were obtained.

### Sphere Migration Assay

To analyze migration using GSC tumor spheres 2,000 to 3,000 cells were seeded into u-shaped 96-well plates 1-2 days prior to the experiments. Additionally, conventional 96-well plates (“flat wells”) were coated using 10 µg/ml Laminin (Sigma-Aldrich) either overnight at 4°C or for 3 h at 37°C. Shortly before the experiments the plates were pre-warmed in case of overnight-coating and laminin was removed. Two wells of the u-shaped 96-well plates were pooled into one well of the coated plate, thereby ensuring that at least one sphere is placed into every well. The plates are immediately placed into an incubator for 30 minutes. During that time the spheres will start to adhere to the plate. Hereafter the treatment medium is added and images of the entire wells are taken using a Tecan Spark plate reader tempered at 37°C (Tecan). After the indicated timepoints additional pictures are acquired. To analyze cellular migration out of the spheres the corresponding pictures are analyzed using FIJI (Schindelin *et al*. 2012). After applying the “find edges” function the pictures are merged in suitable false-color channels. Then, the most migrated cell is manually selected using the line-tool starting from the sphere at the first timepoint (t0) to the timepoint of interest (tx). Starting from this cell 3 additional cells in 90°C angle are measured. By applying the correct pixel/size ratio using the “set scale” function the migration distance in µm was obtained.

### Sphere Invasion Assay

This assay was performed to analyze the invasive potential of GSCs. Corning® Bio-Coat® Matrigel® Invasion Chambers with 8.0 μm PET membranes were rehydrated for 2 h at 37 °C using NB-A medium with only 100 U/ml Penicillin 100µg/mL Streptomycin (Gibco) added. During incubation, the cells were prepared in NB-A medium containing only Penicillin and Streptomycin at a cell density of 80,000 cells per chamber (Hira *et al*. 2020). This supports the invasive behavior of the cells, by generating a growth factor dependent gradient, by placing the chambers in NB-A complete medium (including growth factors). After rehydration of the chambers, the medium was removed and cells were pipetted in the corresponding chambers alongside adding complete medium under the chambers, guaranteeing the growth factor gradient. After this step, the chambers were incubated for 72 h at 37 °C. After incubation, the medium including the seeded cells was removed. Following with a fixation step using 1 ml of methanol (Sigma-Aldrich) incubating for 2 min. After removing the chambers from the methanol, the chambers were stained with 1 ml of 0.1 % crystal violet (Sigma) for 2 min. The chambers were washed with distilled H2O and placed in a new 24-well plate waiting to dry. During the drying process, cells which did not migrate through the Matrigel, were removed using a cotton swab. The chambers were then analyzed with a fluorescent microscope (Nikon eclipse TE2000-S). The invaded cells were counted individually using 7 vision fields and a 20x objective. An overview picture was generated with a 10x objective.

### Adult Organotypic Slice Cultures and Ex Vivo Tumor Growth assay

Adult organotypic tissue slice culture (OTC) was carried out as described previously (Remy *et al*. 2018; Linder *et al*. 2019). Mouse brains were dissected and the dura mater was removed. Subsequently, mouse brains were placed in warm (35-37 °C) 2% low-melting agarose (Carl Roth). Using a VT1000 Vibratome (Leica, Wetzlar, Germany) 150 to 200 µm thick transverse sections were obtained. The sections were transferred to parafilm and kept moist using a drop of PBS. Hereafter the brain slices were cut into pieces of approximately 2 × 5 mm to ensure that they fit into FEP-foil container for subsequent microscopy. These sections were placed on Millicell cell culture inserts (Merck KGaA, Darmstadt, Germany) and cultured in 6-well plates using FCS-free medium consisting of DMEM/F12 supplied with 1x B27, 1x N2 supplement and 100 U/ml Penicillin 100µg/mL Streptomycin (all from Gibco). One day later, one tumor sphere was placed into the center of each brain slice snippet (day 0). Adequate spheres were prepared by seeding 2000 - 3000 cells/well in u-shaped 96-well plates using 200 µl Medium. They were allowed to grow for 1-3 days in order to aggregate into one large sphere. One day after sphere spotting into the brain slices the treatment was started and refreshed 3 times per week. Tumor growth was evaluated after pictures have been taken regularly by Nikon SMZ25 stereomicroscope equipped with a P2-SHR Plan Apo 2 x objective operated by NIS elements software.

conventional microscopy or transferred into PBS and kept at 4°C until LSFM-acquisition.

### Specimen preparation and mounting in the Light Sheet Microscope

#### Specimen holder production and assembly

A new LSFM specimen holder suitable for the size of the brain OTC used in the work was designed with a CAD software (the open-source parametric CAD-software FreeCAD or Fusion360 by Autocad). Next, the model was converted to stereolithographic format (STL) for 3D-printing. 3D printing was performed with an LCD stereolithographic printer (Anycubic Photon S) (Supplementary Figure 1).

**Figure 1:**
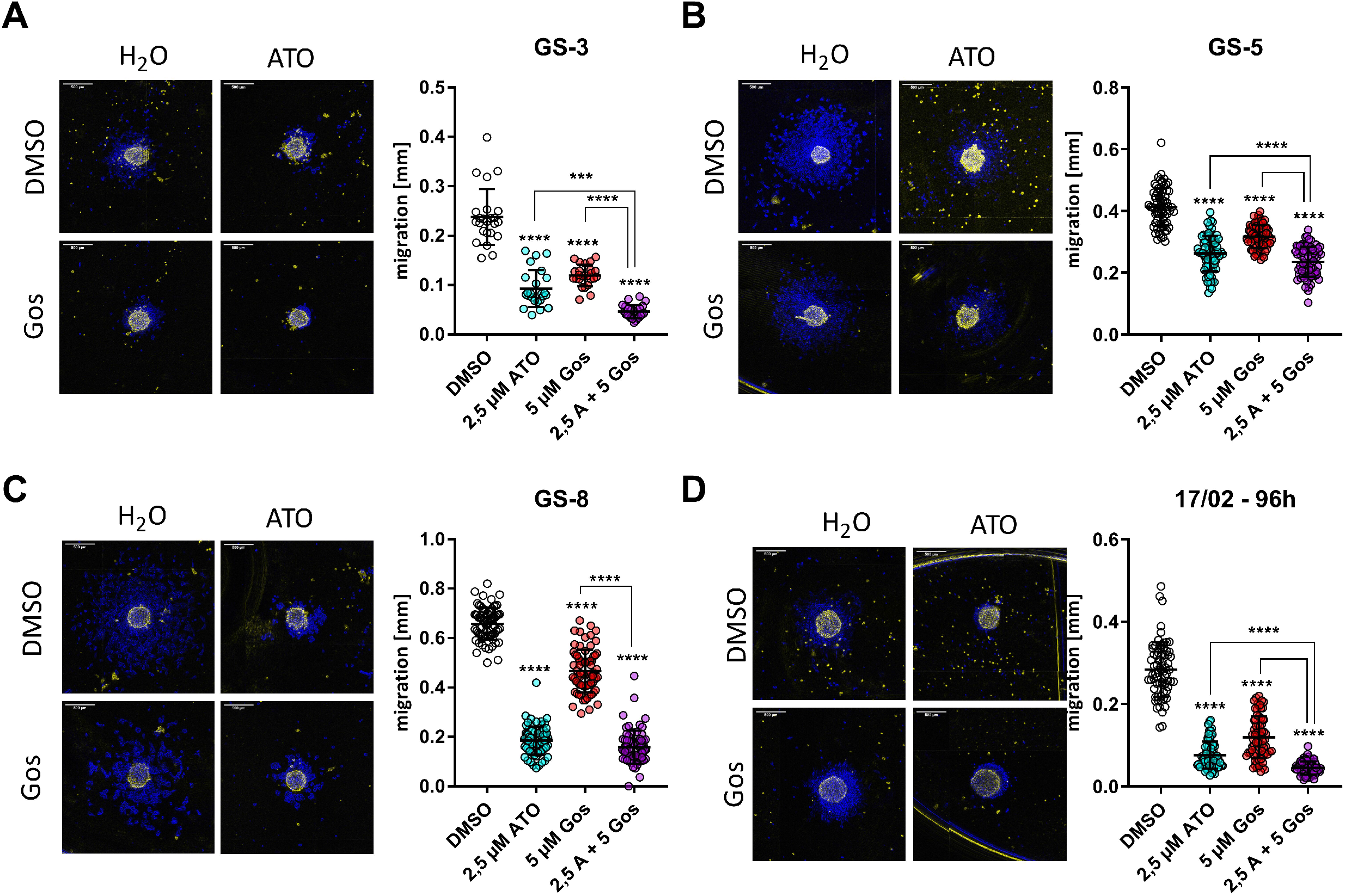
Arsenic trioxide (ATO) alone or in combination with (-)-Gossypol (Gos) effectively blocks migration of GSCs. Sphere Migration assays of (A) GS-3, (B) GS-5, (C) GS-8 and (D) the primary culture 17/02 were treated with sub-lethal concentrations of ATO (2.5 µM) or Gos (5 µM) or solvent (H2O and DMSO, respectively) and imaged 20 min after transferring to laminin-coated 96-well plates (false color, yellow) or 24h and in case of 17/02 96h later (false color, blue) and the migration distance was measured using FIJI (Schindelin et al. 2012). The point-plots show the summary of at least three experiments performed in triplicates. Scale bar: 500 µm. *: p<0.05; **: p<0.01; ***: p<0.001; ****p>0.0001; Two-Way ANOVA with Tukey’s multiple comparison test (GraphPad Prism 9)

FEP-foil custom cuvettes were realized following the procedure described in (Hotte *et al*. 2019). Briefly, 3D-printed positive molds containing a 4×4 array of square pillars with a cross-section of 2mm × 2mm and a length of 4mm were used as vacuum thermoforming tools. After heating a roughly 10cm × 10cm 50 µm-thick FEP foil in the thermoforming machine, vacuum was applied, and the shape of the positive mold impressed in the foil. Next, the individual FEP-foil cuvettes were cut out of the array and glued with UV-sensitive resin (Anycubic) or two-component epoxidic glue onto the top of the holder (supplementary Figure 2).

**Figure 2:**
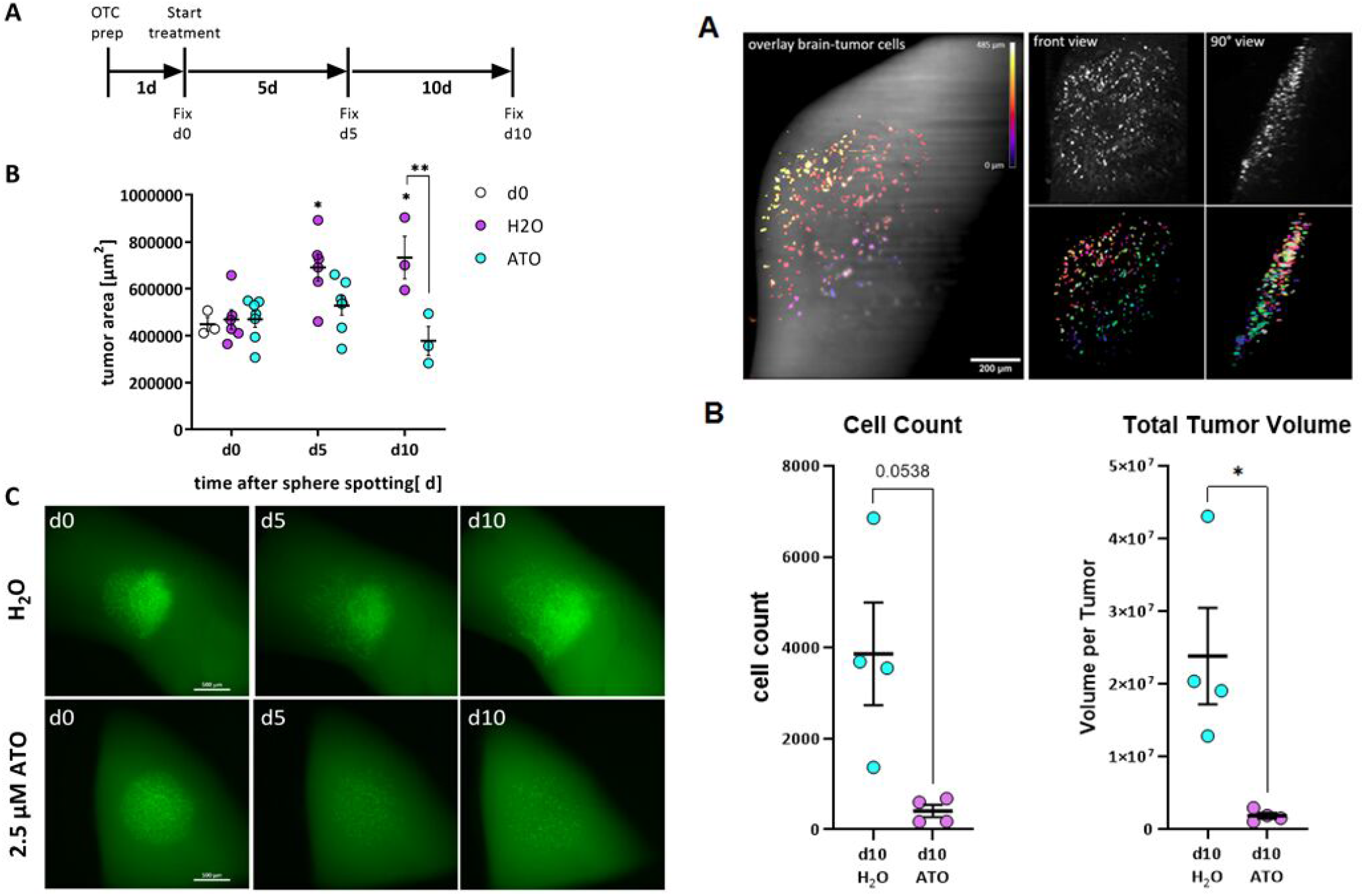
Arsenic Trioxide effectively reduces tumor growth in adult organotypic tissue slice culture based on epifluorescence microscopy and OTCxLSFM-based single cell analysis. (A) Timeframe of the experiments. (B) Quantification of tumor area before treatment (d0) and after 5 (d5) and 10 days (d10) after treatment with 2.5 µM arsenic trioxide (ATO) or solvent (H2O). (C) Microscopic pictures of the same tumor after treatment with solvent (H2O) or ATO. (D). Left panel: overlay of brain slice and tumor cells. The image results from overlaying the maximum projection of the autofluorescence signal from the brain with the GFP signal emitted by the tumor cells. The color coding represents the depth in the image stack. Right panel, top: the front and 90° views of the tumor cells are shown (both maximum projections of the original image stack). Right panel, bottom: front and 90° views of the corresponding segmented and labeled 3D data set. The “Total Tumor Volume” and “cell count” parameters shown in the E, F panels are extracted from the segmented 3D data set. (G and H) Pooled data of both experiments reveal efficient depletion of tumor volume and cell count after ATO-treatment. The values presented in (E and F), as well as supplemental figure 3 were first normalized to H2O-treated tumor per experiment and subsequently pooled to perform statistical analyzes. *: p<0.05; **: p<0.01, Unpaired T-Test with Welch’s Test was performed (GraphPad Prism 9).

#### Mounting the OTC specimen in the holder

Individual PFA-fixated and with CUBIC-2 optically cleared OTC snippets were picked-up from the multi-well plate with a fine-tip brush and carefully deposited in the FEP-foil cuvette glued on the specimen holder (supplementary Figure 2). The cuvette was previously partially filled up with CUBIC-2 solution, to fully embed the OTC specimen in the clearing solution.

#### LSFM imaging

LSFM is a three-dimensional fluorescence microscopy that allow a fast acquisition of image stacks in large and highly scattering samples. Optical sectioning of the sample is achieved through a laser light sheet, which selectively excites fluorescence in single plane of the sample at one time. The emitted light in each plane is collected by an objective lens arranged perpendicular to the optical axis of the light source and detected by a sensitive camera (CCD or CMOS). By axially translating the sample through the light sheet, a three dimensional data set is recorded. An advantage using LSFM is that only the imaged plane is excited by the light sheet. Thus, photobleaching and phototoxicity of the sample are reduced compared to confocal microscopy. Furthermore, the acquisition time is decreased and LSFM provides a new opportunity to investigate the biology of cells and other microorganisms with high spatial and temporal resolution (Pampaloni *et al*. 2015).

For LSFM imaging, a custom-built “monolithic digitally-scanned light sheet microscope” (mDSLM) was used (Pampaloni *et al*. 2013). The mDSLM features a motorized xyz?-stage placed below the specimen chamber. Light sheet imaging was performed with an Epiplan-Neofluar 2.5x/0.06 illumination objective (Carl Zeiss), an N-Achroplan 10x/0.3 detection objective (Carl Zeiss) and a Neo CCD camera (ANDOR Technology, Ireland). Laser wavelength and bandpass filter sets (centre wavelength/full-width at half maximum): 561 nm, 607/70 nm; 488 nm; 525/50 nm, 405 nm, 447/50 nm. Prior to imaging, the mounted OTC specimen was carefully inserted in the mDSLM chamber, also filled with CUBIC-2 solution. After a correct positioning of the specimen to visualize the tumor in its entirety, the image stack recording was started.

### Image processing pipeline

The open-source Fiji (ImageJ version 1.53q, Java version 1.8.0_172, 64 bit) was used to process the raw image stack recorded with the light sheet microscope. The raw data was all saved as uncompressed tif-stacks. **Preprocessing**. After uploading in Fiji, the stacks were first cropped to the region of interest (ROI). Next, the data was denoised with the ImageJ plugin”SNR V2 Variational Stationary Noise Remover” (Escande *et al*. 2017) to suppress stripe artifacts deriving from the light sheet illumination. Thereafter, the background intensity was subtracted from every slice using the “Subtract Background” function (Fiji, ball radius of 50 pixels). Next, a median filter was applied to the image stack. **Segmentation**. For segmentation of the pre-processed data, the Fiji/ImageJ plugins 3D ImageJ Suite (Ollion *et al*. 2013) and MorphoLibJ (Legland *et al*. 2016) were used. First, the seeds corresponding to each cell in the 3D stack were determined with the “3D Maxima Finder” function (3D ImageJ Suite). Next, the “3D Spot Segmentation” function (3D ImageJ Suite) was applied. The resulting segmented 3D data set was post-processed by using MorphoLibJ tools “Labels Size Filtering”, which allows for the removal of segmented debris particles, and “Label Edition”, which allows to manually correct further segmentation artifacts. **Data extraction**. Finally, the 3D cell centroids and the various 3D morphological parameters shown in Figure 3 were extracted by using the 3D ImageJ Suite functions “3D Centroid” and “3D Shape Measure”, respectively.

**Figure 3:**
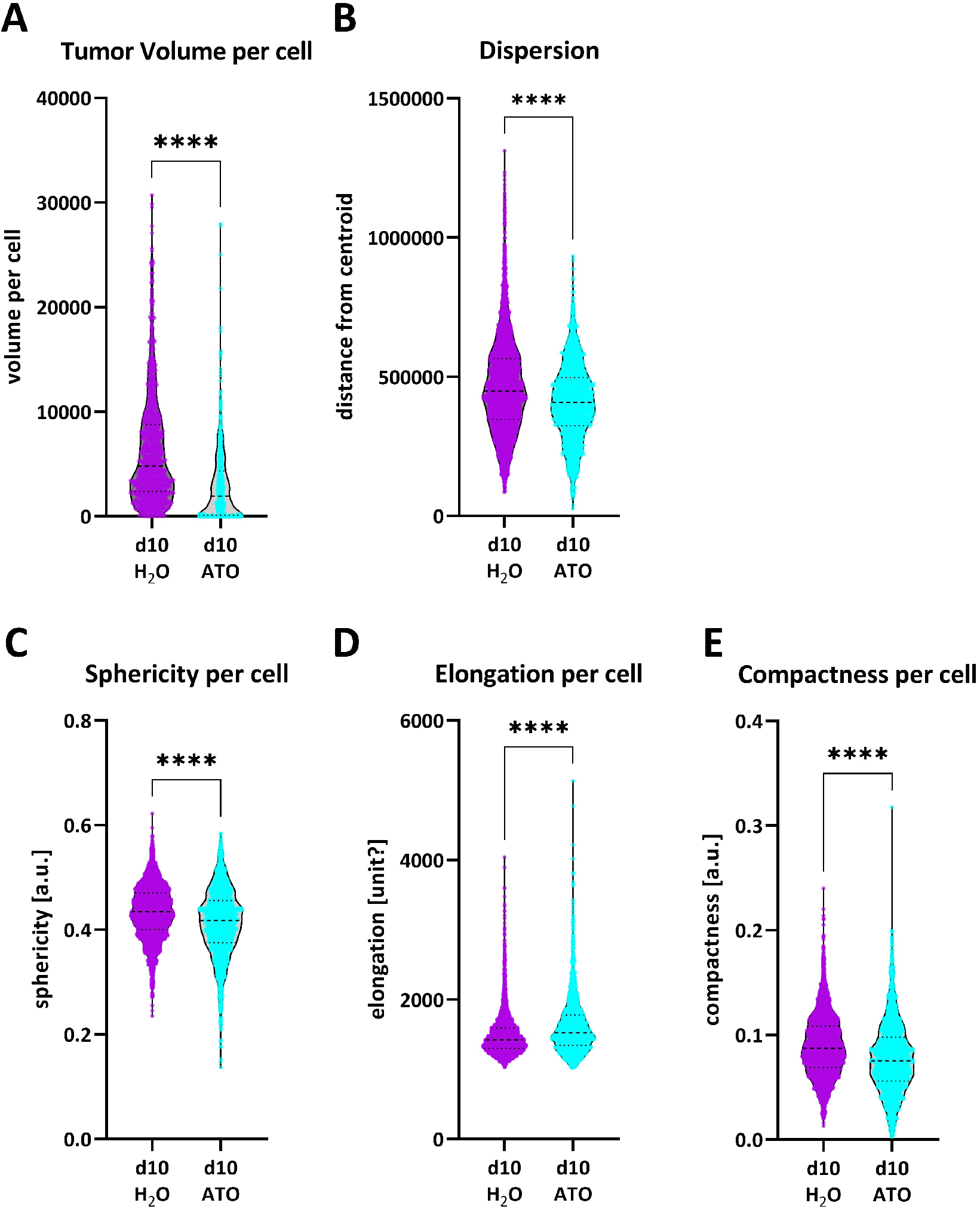
Morphological parameters extracted from the segmented 3D data set. *The parameters are for GSC line GS-5 treated with 2.5µM ATO for (A) the tumor volume per cell, (B) the tumor dispersion, i.e. the distance to the tumor centroid, (C) the sphericity per cell, (D) the elongation per cell and (E) the compactness per cell*. Unpaired T-Test with Mann-Whitney test was performed. Error bars are SEM. * p < 0.05; ** p < 0.01; *** p < 0.001; **** p < 0.0001 against H_2_0 treated cells.

## Results

### Arsenic trioxide inhibits migration in vitro and tissue infiltration ex vivo

Based on our previous work showing that the combination of arsenic trioxide (ATO) + Gossypol (Gos) reduces stemness of GSCs and regulates many proteins that can be clustered under the term “movement” (Linder *et al*. 2019) we hypothesized that ATO (+/-Gos) might be able to inhibit the migratory capacity of GSCs. In order to analyze migration using free-floating GSC spheres we developed an assay that employs the capacity of GSCs to grow adherently on laminin-coating plates, without changing their stem-like phenotype (Pollard *et al*. 2009). Using this assay we could show that within 30 minutes after transferring spheres onto coated plates, the sphere attaches to the plates and the cells start to migrate. Therefore we applied ATO, Gos and the combination of both to three GSC lines, GS-3, GS-5 and GS-8 (Gunther *et al*. 2008) and one primary culture, 17/02, (Linder *et al*. 2019), which are all sensitive to ATO-treatment (Linder *et al*. 2019) (Fig. 1). This experiment revealed that in all 4 cell lines both single treatments effectively block migration, whereas ATO is more effective than Gos. The combination treatment is more effective than ATO-treatment in GS-3, GS5 and 17/02 indicative of the synergism between the two drugs that we previously determined.

In order to apply our newly developed method and to test its capacity for three-dimensional analyses, we performed a proof-of-concept experiment by treating GS-5 GFP/Luc-tumors (Linder *et al*. 2019) using ATO. We chose only ATO-treatment, because it exerts a very strong migration inhibition on its own and we applied GS-5, because in our experience this cell culture has the most consistent growth pattern *ex vivo*. Based on our previous data (Linder *et al*. 2019) we anticipated that after 5 days a reduced tumor growth will occur and that after 10 days a potent reduction in tumor size will be apparent. Accordingly, we chose three timepoints to test the OTCxLSFM method: one day after tumor sphere spotting (treatment d0 “untreated”), 5 days after the initial treatment and 10 days after the initial treatment (Fig. 2 A). For each timepoint we analyzed at least three slices. Firstly, we measured changes in the tumor area over time (see materials and methods and (Remy *et al*. 2018; Linder *et al*. 2019) for each timepoint (Fig. 2 B) and could confirm our previous findings (Linder *et al*. 2019). Before treatment (d0) all tumors are of similar size, whereas after 5 days the solvent-treated tumors are significantly larger, while ATO-treated tumors are not, indicating growth inhibition as anticipated. After 10 days the solvent-treated tumors further increased in size, while ATO-treated tumors are smaller compared to the initial tumor sizes. In fact, the tumors after ATO-treatment are significantly smaller compared to solvent-treatment at the same timepoint. An example for each treatment and timepoint is depicted in figure 2 C. The microscopic evaluation further showed that tumor cells begin to invade the surrounding parenchyma very quickly. This is especially true for solvent-treated tumors (compare H_2_O d5 with d0). In addition, the measurement of the tumor projection area likely underestimates the true effect size of ATO-treatment, since the tumor area does not change significantly the intensity of the GFP-signal, however it is greatly reduced after ATO-treatment, indicating that the tumor cells are effectively depleted. In order to address this discrepancy we applied a three-dimensional analysis of the tumor with light sheet microscopy to obtain a more complete picture.

Figure 2D shows the three-dimensional rendering of a representative organotypic tumor culture (a control specimen treated with solvent) recorded with the light sheet microscope. The GFP fluorescence signal emitted by the tumor cells is nearly one order of magnitude stronger compared to the background autofluorescence emission of the surrounding brain. The background fluorescence is indeed very useful to locate the spreading of the tumor cells in the anatomy of the surrounding brain tissue. The rendered image stack can be rotated and viewed under different angles (a frontal and a lateral view of the tumor are shown in Figure 2G). The migration of the tumor cell into the underlying tissue can be readily visualized by color-coding the depth (Figure 2G). The high signal-to-noise ratio of the fluorescence images allows an efficient segmentation of the individual tumor cells and the quantitative analysis of multiple cell parameters, as shown in Figure 3,5 and 6. The effect of the ATO treatment on tumor volume and cell count, as obtained from the light sheet 3D image stacks is shown in Figure 2E and F. This analysis revealed a strong tendency towards decreased total tumor volume (Fig. 2E) as well as a reduced total cell count per tumor (Fig. 2F). Notably, neither difference reached statistical significance likely due to low sample size (n=3). To address this issue we repeated the experiment and could observe a similar trend (Supplemental Figure 3). Normalizing and pooling the data revealed that both the total tumor volume (Fig. 2G) as well as total cell count (Fig. 2H) are significantly reduced.

**Figure 4:**
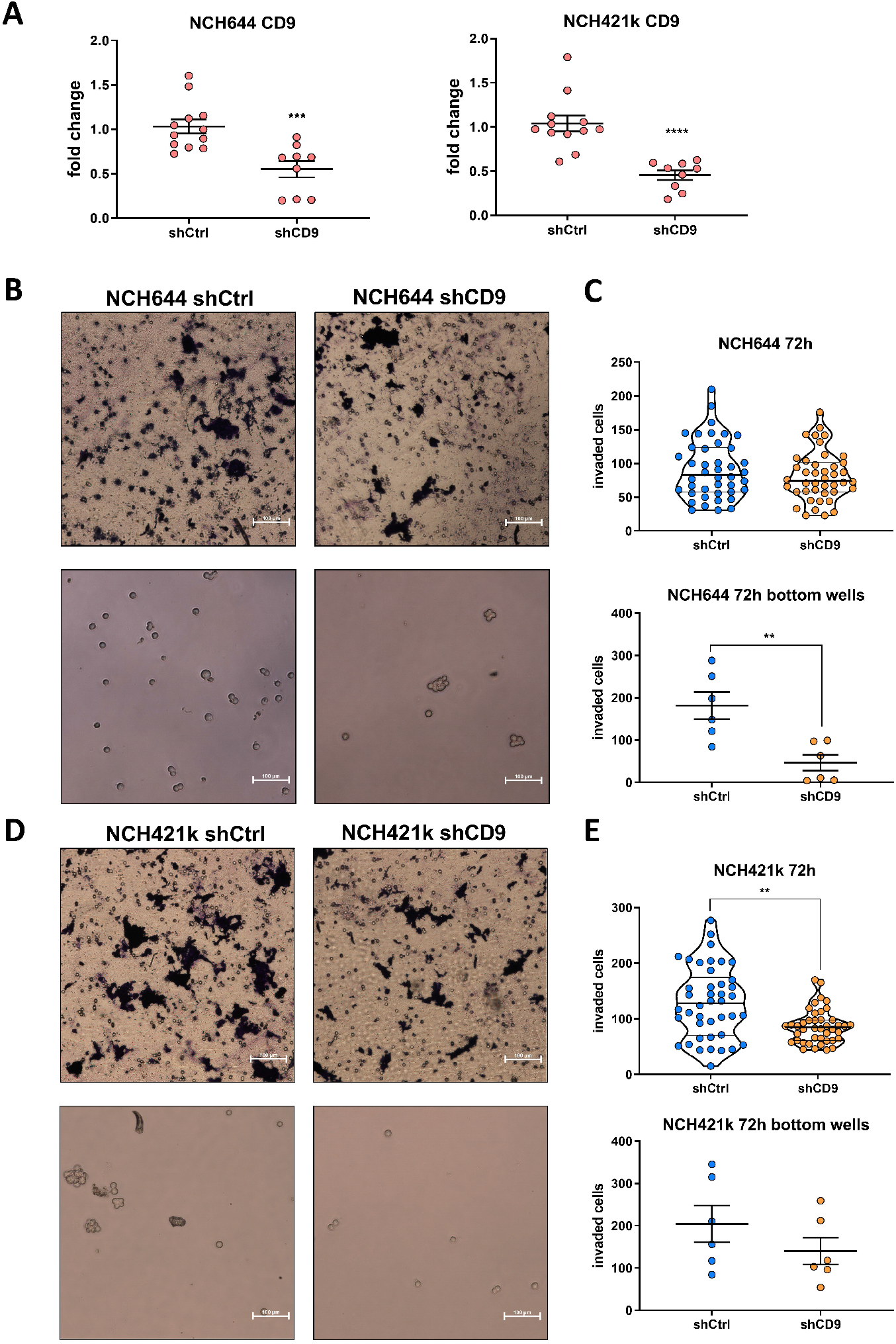
Boyden Chamber invasion assay of NCH644 (A-C) and NCH421k (A;D-E) GSCs after CD9 depletion. (A) Knockdown validation of NCH644 and NCH421k cells show significant decrease of CD9 expression after depletion. Pictures of Boyden Chambers after 72h of incubation generated with using the Nikon eclipse TE2000-S; scale bar 100 μm for (B) NCH644 and (D) NCH421k The quantification is displayed as violin plots in (C) for NCH644 and in (E) for NCH421k. GSCs were seeded once in 3 replicates and afterwards counted in 7 vision fields at 20x. Unpaired T-Test with Mann-Whitney test was performed. Error bars are SEM. * p < 0.05; ** p < 0.01; *** p < 0.001; **** p < 0.0001 against shCtrl cells.

**Figure 5:**
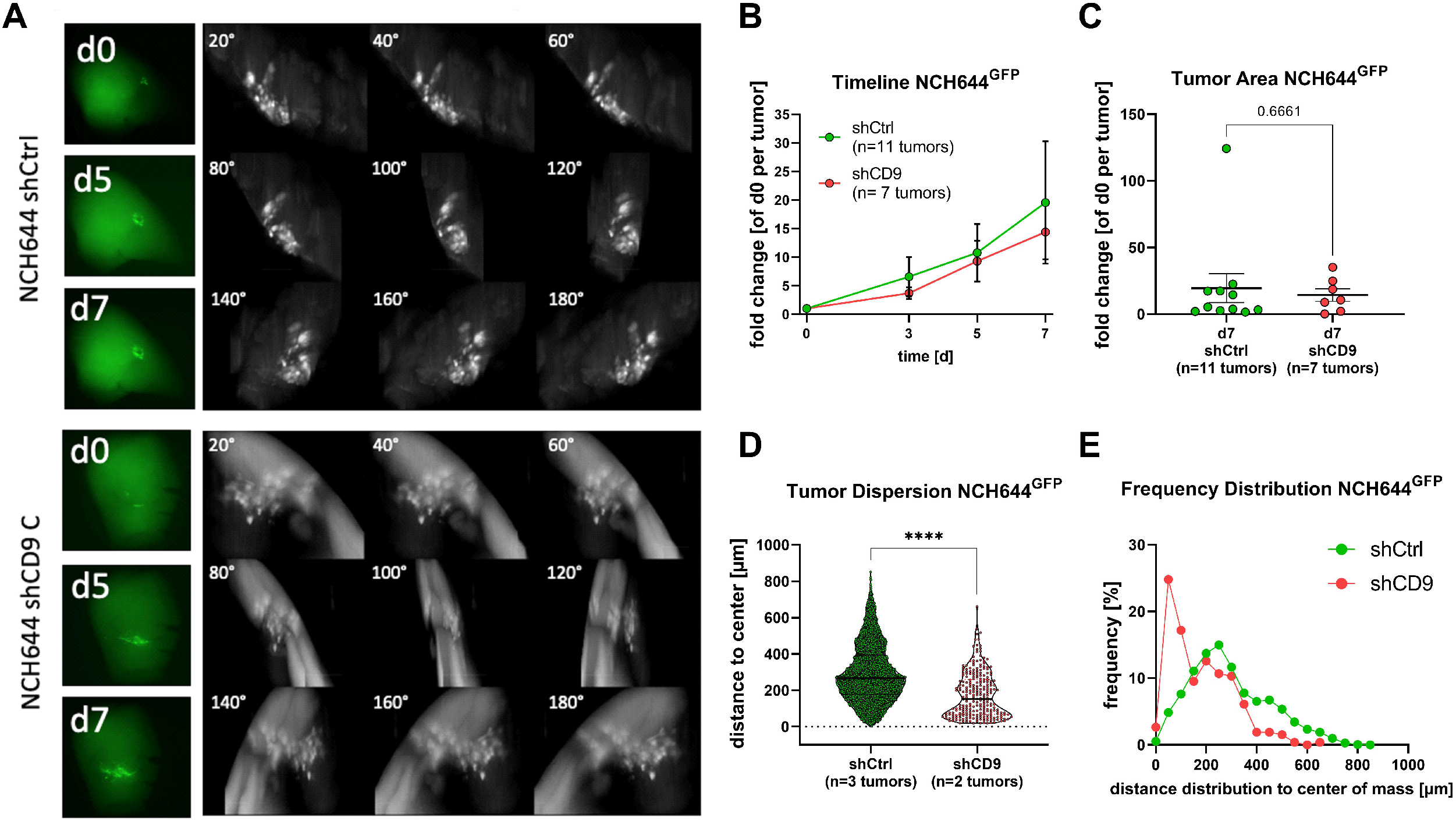
CD9 depletion affects tumor growth in ex vivo organotypic tissue culture tumor growth kinetics in NCH644 cells. (A) Representative microphotographs of tumors transplanted onto adult murine brain organotypic tissue culture slices for NCH644 shCtrl and shCD9 after 0, 5, and 7 d of incubation (left side), as well as montages of 3D-reconstruction of LSFM pictures displayed in 20° angles. (B) Summarized growth kinetic over the entire observation time and (C) point-plot depicting individual fold-changes tumor sizes at the latest timepoint (d7). (D) Violin-Plot depicting tumor cell dispersion, each point represents one tumor cells and (E) Frequency distribution showing the relative amount of migration distance in 50 µm bins. Unpaired T-Test with Mann-Whitney was performed. Error bars are SEM. * p < 0.05; ** p < 0.01; *** p < 0.001; **** p < 0.0001 against shCtrl cells.

**Figure 6:**
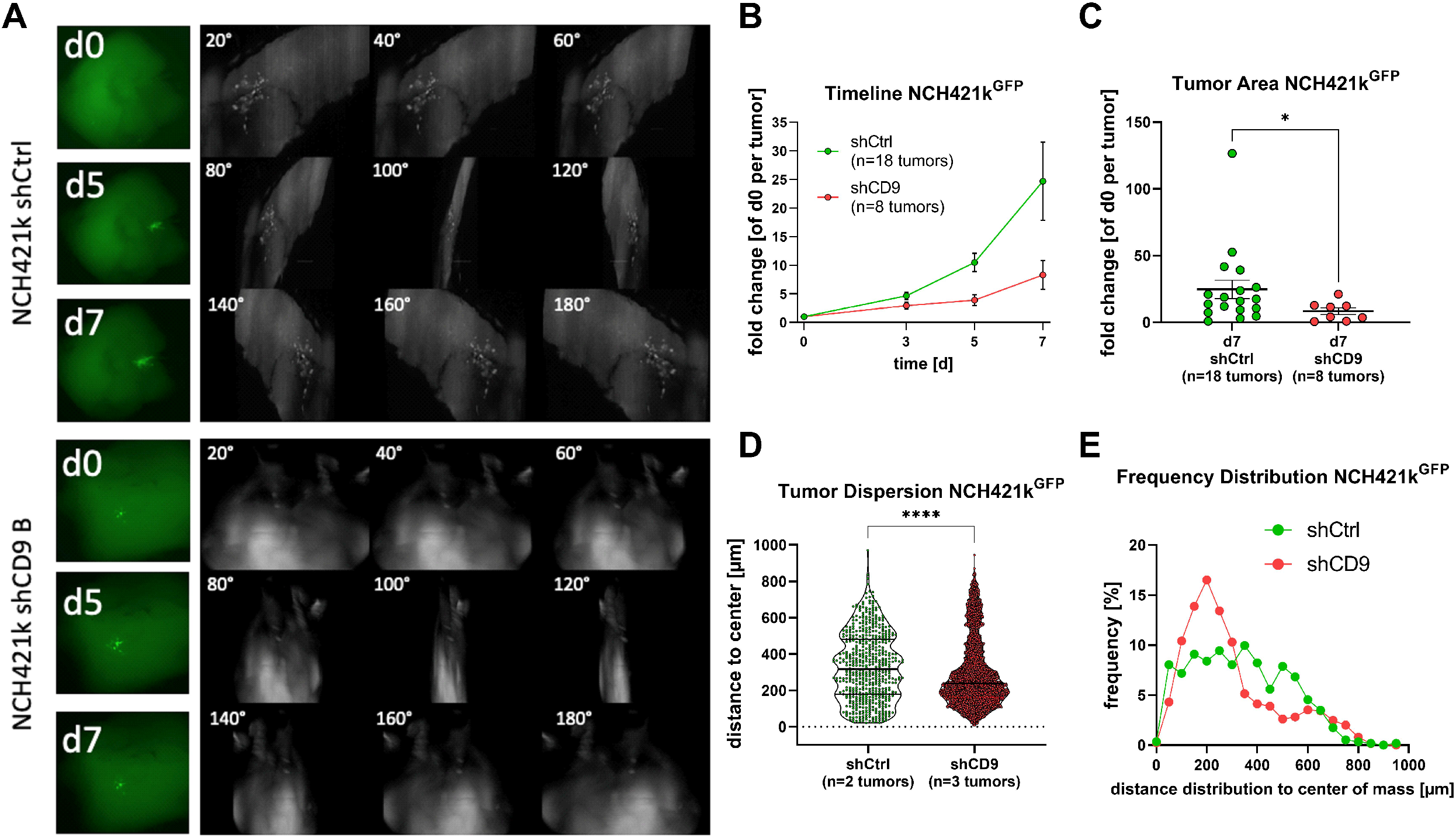
CD9 depletion affects tumor growth in ex vivo organotypic tissue culture tumor growth kinetics in NCH421k cells. (A) Representative microphotographs of tumors transplanted onto adult murine brain organotypic tissue culture slices for NCH644 shCtrl and shCD9 after 0, 5, and 7 d of incubation (left side), as well as montages of 3D-reconstruction of LSFM pictures displayed in 20° angles. (B) Summarized growth kinetic over the entire observation time and (C) point-plot depicting individual fold-changes tumor sizes at the latest timepoint (d7). (D) Violin-Plot depicting tumor cell dispersion, each point represents one tumor cells and (E) Frequency distribution showing the relative amount of migration distance in 50 µm bins. Unpaired T-Test with Mann-Whitney test was performed. Error bars are SEM. * p < 0.05; ** p < 0.01; *** p < 0.001; **** p < 0.0001 against shCtrl cells.

Next, in order to obtain a more complete understanding of the cellular alterations following the ATO treatment we performed a comprehensive cell shape analysis (Figure 3). This method revealed that the overall cellular volume of ATO-treated tumors is prominently and significantly reduced (Fig. 3A). Additionally, we performed an analysis of tumor cell dispersion from the center of the tumor mass (Fig. 3B). This readout is indicative of overall migratory/invasive capacity. As expected from our previous report (Linder *et al*. 2019) and our in vitro data display ATO-treated tumor cells an overall reduced distance from the centroid. In line with these findings, we observed a decrease of cellular sphericity, although this difference was only minor (Fig. 3CE), while the elongation of ATO-treated tumor cells was increased upon treatment (Fig. 3D). Similarly, we observed that the compactness of the cells is diminished (Fig. 3E).

### Stable depletion of CD9 prevents migration in vitro and reduces tissue infiltration ex vivo

Hereafter we wanted to challenge our method by investigating migration and invasion using a genetic knockdown model. We chose to deplete the tetraspanin CD9, because it has been previously shown to be important for GBM migration and invasion (Podergajs *et al*. 2016) and the CD9 protein was among the most decreased hits after ATO+Gos-treatment from our previous report (Linder *et al*. 2019).

For this purpose we first depleted CD9 via stable shRNA-mediated knockdown (CD9-KD) in GS-5 as well as two cell lines that have previously been described to contain high levels of CD9-protein, NCH644 and NCH421k (Podergajs *et al*. 2016). Using GS-5 cells, we successfully confirmed efficient depletion of CD9 protein expression (Sub. Fig. 4A). With these cells we first assessed migration in vitro using the sphere migration assay with GS-5 cells (Sub. Fig. 4B and C). This approach revealed that GS-5 CD9-KD cells migrate slower compared to their parental cell line, showing a significantly lower distance covered after 24, 48 and 72h. After 96h, likely reflecting the maximal migration distance, no more difference is apparent. For NCH644 (Fig. 4A) and NCH421k (Fig. 4B) we could also confirm CD9-depletion via qPCR. Unfortunately, these two cell lines only weakly adhere to laminin-coated surfaces and are therefore not suitable for sphere migration assays. Hence, we performed modified Boyden chamber assay to investigate invasion processes. This approach revealed that for NCH644 (Fig. 4C) almost no difference in the number of invaded cells could be determined for the cells that invaded through the matrix (Fig. 4C upper half), whereas significantly fewer spheres could be determined in the bottom well (Fig. 4C lower half), indicating that NCH644 CD9-KD cells are hindered in their invasive potential. Similarly, NCH421k CD9-KD cells (Fig. 4D) invaded through the matrigel matrix to a reduced extent (Fig. 4D, upper half), which is reflected in fewer spheres in the bottom well (Fig. 4D, lower half). Hereafter, we analyzed changes in stemness marker expression as well as stemness features in both GSCs. This approach revealed a potent reduction of *CXCR4, OLIG2* and *SOX2* expression in both lines (Sup Fig. 5A and B), as well as blockade of sphere forming potential after 7 d of incubation, as shown by fewer and smaller spheres upon CD9-depletion (Sup. Fig. 5C and D).

Hereafter, we performed OTCs with the NCH644 and NCH421k shCtrl and shCD9 cells to analyze if CD9-depletion affects tumor growth and tumor cell infiltration. For NCH644^GFP+^ tumors we observed that over the 7 day observation period CD9-depleted tumors showed a tendency for slower growth compared to CD9-proficient tumors (Fig. 5A and quantification in Fig. 5B). Thus, even at the latest analyzed timepoint (d7, Fig. 5C) only a slight tendency is apparent. However, by applying our analysis pipeline after image acquisition using OTCxLSFM, the tumor cell dispersion (Fig. 5D) is significantly reduced, indicating reduced invasion after CD9-KD. Similarly, the frequency distribution (Fig. 5E) showed that more than 20% tumor cells with CD9-KD only migrate up to 100 µm, and also display a skewed frequency distribution. In contrast, CD9-proficient cells display a more bell-shaped distribution with a mean migration distance of approximately 250 µm. Additionally, CD9-proficient cells migrate up to over 800 µm, whereas CD9-KD cells only migrate to a maximum of approximately 600 µm.

For NCH421k very similar results were obtained (Fig. 6). As depicted in the representative epifluorescence images, as well as the montages of the 3D-reconstruction after OTCxLSFM (Fig. 6A) it is apparent that the NCH421kGFP+ shCD9 tumors are smaller and less invasive. This can also be quantified in the covered area according to the epifluorescence analysis (Fig. 6B). Our three-dimensional approach revealed a reduced tumor cell dispersion (Fig. 6C), indicative of reduced invasiveness, as well as a left-skewed frequency distribution after CD9-depletion (Fig. 6D), similar to NCH644 tumors.

In contrast, by analyzing the same samples using “classical” confocal microscopy (Sup. Fig. 6) we could not confirm our findings obtained in NCH644. Hence, no difference in tumor cell dispersion was found and the frequency distribution showed for both, shCtrl and shCD9-cells, a left-skewed distribution. Notably, the farthest migration distance was 400 µm in both conditions and likely represents a technical limitation. For NCH421k shCtrl and shCD9 cells a decreased tumor cell dispersion could be validated, but, again, the maximum migrated distance was only approximately 450 µm. We therefore concluded that only the acquisition of the entire tumor using OTCxLSFM outcompetes the time-consuming analysis of a fraction of the tumors.

## Discussion

Here we present a novel approach to analyze and quantify GBM migration and invasion in a pathophysiological environment. By combining two state-of-the-art methods, 1) murine, adult organotypic (OTC) brain slices to model GBM growth and 2) advanced fluorescence light-sheet microscopy of cleared tissues (LSFM), we developed OTCxLSFM and provide examples of its application to the quantitative analysis of GBM invasion. One of the major novelty of the OTCxLSFM assay is that it provides access the three-dimensional migratory behaviour of the cells. The three-dimensional data sets obtained with OTCxLSFM allows to avoid biases arising from neglecting the depth in 2D images. In fact, in 2D images the invasion of the brain tissue by the GBM cells cannot be quantified, whereas the ability of tumor cells to spread into the tissue’s extracellular matrix rather than on the surface is a key factor determining the letality of the the tumor and its response to the pharmacological treatment. Another important benefit of the OTCxLSFM assay consists in the much faster imaging speed of LSFM compared to the confocal microscope. This allows to measure >10 specimens in their entirety in one imaging session without compromises in terms of depth acquisition.

The development of suitable, complex preclinical models is a necessity to reflect the complexity of a tumor such as GBM. GBM is characterized by its profound heterogeneity, both between patients (intertumoral), but also within the same individuum (intratumoral) (Robertson *et al*. 2019). Complicating matter further is the diffuse infiltration, as well as tissue penetration, mediated via diverse mechanism such as collective invasion, perineuronal satellitosis, diffuse infiltration as well vessel co-option (Chouleur *et al*. 2020; Seano and Jain 2020). Many approaches exist to tackle this complexity, both with their own advantages and disadvantages (reviewed in (Souberan and Tchoghandjian 2020)) and we present the next-level development of OTCs. The concept to study GBM growth using brain slices has been applied in multiple studies (de Bouard *et al*. 2007; Parker *et al*. 2017; Parker *et al*. 2018) mainly employing post-natal brain slices derived from mice, but also from rats grown under high-serum species-unmatched serum-supplementation (Ren *et al*. 2015; Ghoochani *et al*. 2016; Sidorcenco *et al*. 2020), mainly focusing on tumor growth. Some reports show applicability towards analysis on angiogenesis and tumor microenvironment however, these studies mainly rely on further sectioning the slices thereby further increasing the workload. More recent studies, including our own, developed an OTC platform based on adult murine brain slices grown under serum-free conditions (Marques-Torrejon *et al*. 2018; Meyer *et al*. 2018; Remy *et al*. 2018; Linder *et al*. 2019; Gerstmeier *et al*. 2021), thereby ensuring that tumors grow in a more relevant environment, because GBM is a tumor of adult patients. One major drawback of current approaches is the lack of properly investigating the three-dimensional tissue invasion of GBM cells within these slices. Although this problem can be addressed by further performing fine sections of these slices or by employing tissue-clearing and confocal-microscopy, these approaches are either very labor-intensive and time-consuming or, in the case of confocal microscopy, usually restricted to a small fraction of the entire tumor. In contrast, with LSFM it is possible to image very large specimens, while maintaining single cell resolution, thereby greatly reducing the needed acquisition time by simultaneously increasing the possible sample size.

As a proof-of-concept approach we chose ATO as a pharmacological agent and CD9 for the genetic model. We have recently shown that the combination of ATO and Gos can effectively block GSC stemness and a proteomic analysis revealed that almost 40% of all decreased proteins were related to movement (Linder *et al*. 2019) and included CD9 as one the most depleted proteins (Linder *et al*. 2019). We confirmed these data using 4 GSCs lines showing that in particular ATO and the combination of ATO and Gos can block migration. Since ATO alone was already highly effective we chose to only apply the ATO single-treatment. Using an OTC-approach we first confirmed our previous results (Linder *et al*. 2019), that ATO leads to continuous reduction of tumor area and can, after 10 days of treatment, also reduce the area below the original size. An analysis using OTCxLSFM revealed that not only the area on-top of the slices is reduced, but also tissue invasion is greatly disturbed while inducing morphological changes of the cells indicative of reduced invasiveness. Next, we wanted to show the applicability of our novel tool using a transgenic approach. For this purpose we depleted CD9 in GS-5, as well as in two additional cell lines NCH644 and NCH421k (Campos *et al*. 2014), which have been previously shown to express high levels of CD9 and displayed reduced migration/invasion after CD9-depletion (Podergajs *et al*. 2016). After confirmation of reduced migration potential in all three GSC lines we next performed OTCs of NCH644 and NCH421k GSCs and showed that NCH644 cells do not display reduced tumor area, which is in contrast to the previously described improved survival of tumor-bearing mice (Podergajs *et al*. 2016), while NCH421k displayed smaller tumors in line with the literature (Podergajs *et al*. 2016). However, after quantification of the light sheet images, it is proven that both GSCs showed reduced migration of tumor cells into the surrounding brain tissue. In contrast, by only analyzing a subset of the entire tumor using confocal microscopy, we could only validate these results in one of the two cell lines. This demonstrates that it is an absolute necessity to analyze the complete tumor including distant migration of individual tumor cells. The analysis of a randomly selected subset brings in an unnecessary bias, which will lead to false interpretation. In addition, the acquisition via OTCxLSFM is much faster, which is also an important factor, when considering to increase the throughput. One notable advantage of confocal microscopy versus LSFM is the improved image quality and lack of artifacts that frequently occur using light sheet approaches, which eases the sample processing.

In summary, we present a novel analysis tool by combining two state-of-the-art techniques; adult brain slices to assay tumor growth and multifluorescent LSFM to analyze large and complex samples on a single cell level in 3D. We anticipate that this system will provide novel valuable insights into GBM migration/invasion processes and that our analytical pipelines are transferable to other complex model systems.

## Conflict of interest

The authors declare that they have no conflict of interest.

## Author contributions

Conzeptualization: FP,BL; Data curation: AH, FP, BL; Formal analysis: AH, AW, FP, BL; Funding acquisition: FP, BL; Investigation: AH, AW, FP, BL; Methodology: AH, DK, FP, BL; Project administration: BL; Resources: CHM, DK, FP, BL; Software: AH, FP; Supervision: DK, FP, BL; Validation: AH, BL; Visualization: AH, AW, FP, BL; Writing - Original draft: AH, FP, BL; Writing – review & editing: AH, BL

## Acknowledgements

The authors would like to thank Hildegard König for ongoing excellent technical assistance.

## Funding

This study was supported by the Deutsche Forschungsgemeinschaft (DFG; German Research Council) to BL (LI 3687/2-1). Research in the group of FP is supported by EU FET-Open (828931), EU Transition Open (01 101057894), Wilhelm Sander-Stiftung (2020.008.1) and the German Aerospace Center (DLR) (IMMUNO3D-SHAPE (grant number 50WB2019)

## Supplementary Figure Legends

***Supplementary Figure 1*** *(A) 3D rendering of the CAD design of the thermoforming positive molds for the LSFM holders. An array of 8 positive molds is shown. The molds have a square 2 mm × 2 mm cross-section and a height of 5 mm. (B) the real 3D-printed molds array. (C) one thermoformed FEP-foil LSFM holder glued to ist support. (D) a set of ready-to-use LSFM holders*.

***Supplementary Figure 2*** *Specimen mounting. (A) one brain OTC is placed in the holder by gently displacing it with a fine-tip paint brush. (B) appearance of three OTC inside the FEP-foil specimen holder. (C) The specimen holder with the OTC is subsequently filled with the CUBIC-2 solution*.

***Supplemental Figure 3: Arsenic Trioxide reduces tumor growth in the validation experiment using adult organotypic tissue slice culture based on epifluorescence microscopy and OTCxLSFM-based single cell analysis***. *(A). The “Total Tumor Volume” and (B) “cell count” parameters were extracted from segmented 3D data sets of adult OTCs treated for 10 days with 2.5 µM ATO or solvent (H2O) and image acquisition and processing using OTCxLSFM. *: p<0.05; **: p<0.01, Unpaired T-Test with Welch’s Test was performed (GraphPad Prism 9)*.

***Supplemental Figure 4: CD9 depletion leads to reduced migration in GS-5 GSCs***. *(A) The expression of CD9 in GS-5 GSCs after lentiviral transduction of an shCD9 plasmid was analyzed via Western Blot and quantified in relation to GAPDH expression. (B) False color images (generated after 48 h of incubation) are displayed for GS-5 and GS-5 shCD9 GSCs; scale bar 100 μm. (C) GS-5 and shCD9 GSCs showed difference in migration upon CD9 depletion. The experiment was performed three times in 6 technical replicates. Unpaired T-Test with Mann-Whitney test was performed. Error bars are SEM. * p < 0.05; ** p < 0.01; *** p < 0.001; **** p < 0.0001 against GS-5 par*.

***Supplementary Figure 5: CD9-depletion reduces stemness traits in NCH644 and NCH421k GSCs***. *(A and B) Point-plots of Taqman-based qRT-PCR of stemness marker CXCR4, OLIG2 and SOX2 in (A) NCH644 and (B) NCH421k GSCs after stable transduction with shCtrl or shCD9. The experiment was performed three times in biological triplicates. (C and D) Violin plots of the sphere formation assay of (C) NCH644 and (D) NCH421k GSCs. Freshly dissociated GSCs were incubated for 7 d. The experiment was performed three times using 5-10 biological replicates. The mean sphere number/well and the mean sphere area/well was determined for each cell line. One-Way ANOVA with Tukey’s multiple comparisons test was performed. Error bars are SEM. * p < 0.05; ** p < 0.01; *** p <0.001; **** p < 0.0001 against shCtrl*.

***Supplementary Figure 6 Quantification of confocal generated pictures observing CD9 depletion performing ex vivo organotypic tissue culture using GSC line NCH644 and NCH421k shCtrl and shCD9***. *(A) Montages of 3D-reconstruction of confocal pictures displayed in 20° angles for NCH644 GSCs. (B) Montages of 3D-reconstruction of confocal LSFM pictures displayed in 20° angles for NCH421k GSCs. (C) Violin-Plot depicting tumor cell dispersion, each point represents one tumor cell for NCH644 shCtrl (7 tumors) and shCD9 (3 tumors). (D) Frequency distribution showing the relative amount of migration distance in 25 µm bins of NCH644 tumors. (E) Violin-Plot depicting tumor cell dispersion, each point represents one tumor cell for NCH421k shCtrl (9 tumors) and shCD9 (3 tumors). (F) Frequency distribution showing the relative amount of migration distance in 25 µm bins of NCH421k tumors.Unpaired T-Test with Mann-Whitney test was performed. Error bars are SEM. * p < 0.05; ** p < 0.01; *** p < 0.001; **** p < 0.0001 against shCtrl cells*.

## References

Ariza, A., Lopez, D., Mate, J. L., Isamat, M., Musulen, E., Pujol, M., Ley, A. and Navas-Palacios, J. J. (1995). “Role of CD44 in the invasiveness of glioblastoma multiforme and the noninvasiveness of meningioma: an immunohistochemistry study.” Hum Pathol 26(10): 1144–1147.

Becker, K. P. and Yu, J. (2012). “Status quo--standard-of-care medical and radiation therapy for glioblastoma.” Cancer journal 18(1): 12–19.

Boumahdi, S., Driessens, G., Lapouge, G., Rorive, S., Nassar, D., Le Mercier, M., Delatte, B., Caauwe, A., Lenglez, S., Nkusi, E., Brohee, S., Salmon, I., Dubois, C., del Marmol, V., Fuks, F., Beck, B. and Blanpain, C. (2014). “SOX2 controls tumour initiation and cancer stem-cell functions in squamous-cell carcinoma.” Nature 511(7508): 246–250.

Bradshaw, A., Wickremsekera, A., Tan, S. T., Peng, L., Davis, P. F. and Itinteang, T. (2016). “Cancer Stem Cell Hierarchy in Glioblastoma Multiforme.” Frontiers in surgery 3: 21.

Campos, B., Gal, Z., Baader, A., Schneider, T., Sliwinski, C., Gassel, K., Bageritz, J., Grabe, N., von Deimling, A., Beckhove, P., Mogler, C., Goidts, V., Unterberg, A., Eckstein, V. and Herold-Mende, C. (2014). “Aberrant self-renewal and quiescence contribute to the aggressiveness of glioblastoma.” J Pathol 234(1): 23–33.

Campos, B., Wan, F., Farhadi, M., Ernst, A., Zeppernick, F., Tagscherer, K. E., Ahmadi, R., Lohr, J., Dictus, C., Gdynia, G., Combs, S. E., Goidts, V., Helmke, B. M., Eckstein, V., Roth, W., Beckhove, P., Lichter, P., Unterberg, A., Radlwimmer, B. and Herold-Mende, C. (2010). “Differentiation therapy exerts antitumor effects on stem-like glioma cells.” Clinical cancer research : an official journal of the American Association for Cancer Research 16(10): 2715–2728.

Chouleur, T., Tremblay, M. L. and Bikfalvi, A. (2020). “Mechanisms of invasion in glioblastoma.” Current opinion in oncology 32(6): 631–639.

de Bouard, S., Herlin, P., Christensen, J. G., Lemoisson, E., Gauduchon, P., Raymond, E. and Guillamo, J. S. (2007). “Antiangiogenic and anti-invasive effects of sunitinib on experimental human glioblastoma.” Neuro-oncology 9(4): 412–423.

de Gooijer, M. C., Guillen Navarro, M., Bernards, R., Wurdinger, T. and van Tellingen, O. (2018). “An Experimenter’s Guide to Glioblastoma Invasion Pathways.” Trends Mol Med 24(9): 763–780.

Escande, P., Weiss, P. and Zhang, W. X. (2017). “A Variational Model for Multiplicative Structured Noise Removal.” Journal of Mathematical Imaging and Vision 57(1): 43–55.

Garros-Regulez, L., Aldaz, P., Arrizabalaga, O., Moncho-Amor, V., Carrasco-Garcia, E., Manterola, L., Moreno-Cugnon, L., Barrena, C., Villanua, J., Ruiz, I., Pollard, S., Lovell-Badge, R., Sampron, N., Garcia, I. and Matheu, A. (2016). “mTOR inhibition decreases SOX2-SOX9 mediated glioma stem cell activity and temozolomide resistance.” Expert Opin Ther Targets 20(4): 393–405.

Gerstmeier, J., Possmayer, A. L., Bozkurt, S., Hoffmann, M. E., Dikic, I., Herold-Mende, C., Burger, M. C., Munch, C., Kogel, D. and Linder, B. (2021). “Calcitriol Promotes Differentiation of Glioma Stem-Like Cells and Increases Their Susceptibility to Temozolomide.” Cancers (Basel) 13(14).

Ghoochani, A., Yakubov, E., Sehm, T., Fan, Z., Hock, S., Buchfelder, M., Eyupoglu, I. Y. and Savaskan, N. E. (2016). “A versatile ex vivo technique for assaying tumor angiogenesis and microglia in the brain.” Oncotarget 7(2): 1838–1853.

Gilbertson, R. J. and Rich, J. N. (2007). “Making a tumour’s bed: glioblastoma stem cells and the vascular niche.” Nature reviews. Cancer 7(10): 733–736.

Glaser, A. K., Bishop, K. W., Barner, L. A., Susaki, E. A., Kubota, S. I., Gao, G., Serafin, R. B., Balaram, P., Turschak, E., Nicovich, P. R., Lai, H., Lucas, L. A. G., Yi, Y., Nichols, E. K., Huang, H., Reder, N. P., Wilson, J. J., Sivakumar, R., Shamskhou, E., Stoltzfus, C. R., Wei, X., Hempton, A. K., Pende, M., Murawala, P., Dodt, H. U., Imaizumi, T., Shendure, J., Beliveau, B. J., Gerner, M. Y., Xin, L., Zhao, H., True, L. D., Reid, R. C., Chandrashekar, J., Ueda, H. R., Svoboda, K. and Liu, J. T. C. (2022). “A hybrid open-top light-sheet microscope for versatile multi-scale imaging of cleared tissues.” Nature methods 19(5): 613–619.

Gunther, H. S., Schmidt, N. O., Phillips, H. S., Kemming, D., Kharbanda, S., Soriano, R., Modrusan, Z., Meissner, H., Westphal, M. and Lamszus, K. (2008). “Glioblastoma-derived stem cell-enriched cultures form distinct subgroups according to molecular and phenotypic criteria.” Oncogene 27(20): 2897–2909.

Hide, T., Shibahara, I. and Kumabe, T. (2019). “Novel concept of the border niche: glioblastoma cells use oligodendrocytes progenitor cells (GAOs) and microglia to acquire stem cell-like features.” Brain tumor pathology 36(2): 63–73.

Hira, V. V., Breznik, B., Van Noorden, C. J., Lah, T. and Molenaar, R. J. (2020). “2D and 3D in vitro assays to quantify the invasive behavior of glioblastoma stem cells in response to SDF-1alpha.” Biotechniques 69(5): 339–346.

Hof, L., Moreth, T., Koch, M., Liebisch, T., Kurtz, M., Tarnick, J., Lissek, S. M., Verstegen, M. M. A., van der Laan, L. J. W., Huch, M., Matthaus, F., Stelzer, E. H. K. and Pampaloni, F. (2021). “Long-term live imaging and multiscale analysis identify heterogeneity and core principles of epithelial organoid morphogenesis.” BMC Biol 19(1): 37.

Hotte, K., Koch, M., Hof, L., Tuppi, M., Moreth, T., Verstegen, M. M. A., van der Laan, L. J. W., Stelzer, E. H. K. and Pampaloni, F. (2019). “Ultra-thin fluorocarbon foils optimise multiscale imaging of three-dimensional native and optically cleared specimens.” Scientific reports 9(1): 17292.

Krusche, B., Ottone, C., Clements, M. P., Johnstone, E. R., Goetsch, K., Lieven, H., Mota, S. G., Singh, P., Khadayate, S., Ashraf, A., Davies, T., Pollard, S. M., De Paola, V., Roncaroli, F., Martinez-Torrecuadrada, J., Bertone, P. and Parrinello, S. (2016). “EphrinB2 drives perivascular invasion and proliferation of glioblastoma stem-like cells.” Elife 5.

Legland, D., Arganda-Carreras, I. and Andrey, P. (2016). “MorphoLibJ: integrated library and plugins for mathematical morphology with ImageJ.” Bioinformatics 32(22): 3532–3534.

Linder, B., Wehle, A., Hehlgans, S., Bonn, F., Dikic, I., Rodel, F., Seifert, V. and Kogel, D. (2019). “Arsenic Trioxide and (-)-Gossypol Synergistically Target Glioma Stem-Like Cells via Inhibition of Hedgehog and Notch Signaling.” Cancers (Basel) 11(3).

Linder, B., Zoldakova, M., Kornyei, Z., Kohler, L. H. F., Seibt, S., Menger, D., Wetzel, A., Madarasz, E., Schobert, R., Kogel, D. and Biersack, B. (2022). “Antitumor Effects of a New Retinoate of the Fungal Cytotoxin Illudin M in Brain Tumor Models.” International journal of molecular sciences 23(16).

Louis, D. N., Perry, A., Wesseling, P., Brat, D. J., Cree, I. A., Figarella-Branger, D., Hawkins, C., Ng, H. K., Pfister, S. M., Reifenberger, G., Soffietti, R., von Deimling, A. and Ellison, D. W. (2021). “The 2021 WHO Classification of Tumors of the Central Nervous System: a summary.” Neuro-oncology 23(8): 1231–1251.

Luwor, R. B., Baradaran, B., Taylor, L. E., Iaria, J., Nheu, T. V., Amiry, N., Hovens, C. M., Wang, B., Kaye, A. H. and Zhu, H. J. (2013). “Targeting Stat3 and Smad7 to restore TGF-beta cytostatic regulation of tumor cells in vitro and in vivo.” Oncogene 32(19): 2433–2441.

Marques-Torrejon, M. A., Gangoso, E. and Pollard, S. M. (2018). “Modelling glioblastoma tumour-host cell interactions using adult brain organotypic slice co-culture.” Disease models & mechanisms 11(2).

Meyer, N., Zielke, S., Michaelis, J. B., Linder, B., Warnsmann, V., Rakel, S., Osiewacz, H. D., Fulda, S., Mittelbronn, M., Munch, C., Behrends, C. and Kogel, D. (2018). “AT 101 induces early mitochondrial dysfunction and HMOX1 (heme oxygenase 1) to trigger mitophagic cell death in glioma cells.” Autophagy 14(10): 1693–1709.

Ollion, J., Cochennec, J., Loll, F., Escude, C. and Boudier, T. (2013). “TANGO: a generic tool for high-throughput 3D image analysis for studying nuclear organization.” Bioinformatics 29(14): 1840–1841.

Pampaloni, F., Ansari, N. and Stelzer, E. H. (2013). “High-resolution deep imaging of live cellular spheroids with light-sheet-based fluorescence microscopy.” Cell and Tissue Research 352(1): 161–177.

Pampaloni, F., Chang, B. J. and Stelzer, E. H. (2015). “Light sheet-based fluorescence microscopy (LSFM) for the quantitative imaging of cells and tissues.” Cell and Tissue Research 360(1): 129–141.

Parker, J. J., Canoll, P., Niswander, L., Kleinschmidt-DeMasters, B. K., Foshay, K. and Waziri, A. (2018). “Intratumoral heterogeneity of endogenous tumor cell invasive behavior in human glioblastoma.” Scientific reports 8(1): 18002.

Parker, J. J., Lizarraga, M., Waziri, A. and Foshay, K. M. (2017). “A Human Glioblastoma Organotypic Slice Culture Model for Study of Tumor Cell Migration and Patient-specific Effects of Anti-Invasive Drugs.” Journal of visualized experiments : JoVE(125).

Podergajs, N., Motaln, H., Rajcevic, U., Verbovsek, U., Korsic, M., Obad, N., Espedal, H., Vittori, M., Herold-Mende, C., Miletic, H., Bjerkvig, R. and Turnsek, T. L. (2016). “Transmembrane protein CD9 is glioblastoma biomarker, relevant for maintenance of glioblastoma stem cells.” Oncotarget 7(1): 593–609.

Pollard, S. M., Yoshikawa, K., Clarke, I. D., Danovi, D., Stricker, S., Russell, R., Bayani, J., Head, R., Lee, M., Bernstein, M., Squire, J. A., Smith, A. and Dirks, P. (2009). “Glioma stem cell lines expanded in adherent culture have tumor-specific phenotypes and are suitable for chemical and genetic screens.” Cell stem cell 4(6): 568–580.

Quaresma, M., Coleman, M. P. and Rachet, B. (2015). “40-year trends in an index of survival for all cancers combined and survival adjusted for age and sex for each cancer in England and Wales, 1971-2011: a population-based study.” Lancet 385(9974): 1206–1218.

Remy, J., Linder, B., Weirauch, U., Konovalova, J., Marschalek, R., Aigner, A. and Kogel, D. (2018). “Inhibition of PIM1 blocks the autophagic flux to sensitize glioblastoma cells to ABT-737-induced apoptosis.” Biochim Biophys Acta Mol Cell Res.

Ren, B., Yu, S., Chen, C., Wang, L., Liu, Z., Wu, Q., Wang, L., Zhao, K. and Yang, X. (2015). “Invasion and anti-invasion research of glioma cells in an improved model of organotypic brain slice culture.” Tumori 101(4): 390–397.

Richardson, D. S. and Lichtman, J. W. (2015). “Clarifying Tissue Clearing.” Cell 162(2): 246–257.

Robertson, F. L., Marques-Torrejon, M. A., Morrison, G. M. and Pollard, S. M. (2019). “Experimental models and tools to tackle glioblastoma.” Disease models & mechanisms 12(9).

Rominiyi, O., Vanderlinden, A., Clenton, S. J., Bridgewater, C., Al-Tamimi, Y. and Collis, S. J. (2020). “Tumour treating fields therapy for glioblastoma: current advances and future directions.” British journal of cancer.

Schindelin, J., Arganda-Carreras, I., Frise, E., Kaynig, V., Longair, M., Pietzsch, T., Preibisch, S., Rueden, C., Saalfeld, S., Schmid, B., Tinevez, J. Y., White, D. J., Hartenstein, V., Eliceiri, K., Tomancak, P. and Cardona, A. (2012). “Fiji: an open-source platform for biological-image analysis.” Nature methods 9(7): 676–682.

Seano, G. and Jain, R. K. (2020). “Vessel co-option in glioblastoma: emerging insights and opportunities.” Angiogenesis 23(1): 9–16.

Sidorcenco, V., Krahnen, L., Schulz, M., Remy, J., Kogel, D., Temme, A., Krugel, U., Franke, H. and Aigner, A. (2020). “Glioblastoma Tissue Slice Tandem-Cultures for Quantitative Evaluation of Inhibitory Effects on Invasion and Growth.” Cancers (Basel) 12(9).

Soeda, A., Park, M., Lee, D., Mintz, A., Androutsellis-Theotokis, A., McKay, R. D., Engh, J., Iwama, T., Kunisada, T., Kassam, A. B., Pollack, I. F. and Park, D. M. (2009). “Hypoxia promotes expansion of the CD133-positive glioma stem cells through activation of HIF-1alpha.” Oncogene 28(45): 3949–3959.

Souberan, A. and Tchoghandjian, A. (2020). “Practical Review on Preclinical Human 3D Glioblastoma Models: Advances and Challenges for Clinical Translation.” Cancers (Basel) 12(9).

Stelzer, E. H. K., Strobl, F., Chang, B.-J., Preusser, F., Preibisch, S., McDole, K. and Fiolka, R. (2021). “Light sheet fluorescence microscopy.” Nature Reviews Methods Primers 1(1): 73.

Stupp, R., Hegi, M. E., Gilbert, M. R. and Chakravarti, A. (2007). “Chemoradiotherapy in malignant glioma: standard of care and future directions.” Journal of clinical oncology : official journal of the American Society of Clinical Oncology 25(26): 4127–4136.

Stupp, R., Hegi, M. E., Mason, W. P., van den Bent, M. J., Taphoorn, M. J., Janzer, R. C., Ludwin, S. K., Allgeier, A., Fisher, B., Belanger, K., Hau, P., Brandes, A. A., Gijtenbeek, J., Marosi, C., Vecht, C. J., Mokhtari, K., Wesseling, P., Villa, S., Eisenhauer, E., Gorlia, T., Weller, M., Lacombe, D., Cairncross, J. G. and Mirimanoff, R. O. (2009). “Effects of radiotherapy with concomitant and adjuvant temozolomide versus radiotherapy alone on survival in glioblastoma in a randomised phase III study: 5-year analysis of the EORTC-NCIC trial.” The Lancet. Oncology 10(5): 459–466.

Stupp, R., Mason, W. P., van den Bent, M. J., Weller, M., Fisher, B., Taphoorn, M. J., Belanger, K., Brandes, A. A., Marosi, C., Bogdahn, U., Curschmann, J., Janzer, R. C., Ludwin, S. K., Gorlia, T., Allgeier, A., Lacombe, D., Cairncross, J. G., Eisenhauer, E. and Mirimanoff, R. O. (2005). “Radiotherapy plus concomitant and adjuvant temozolomide for glioblastoma.” The New England journal of medicine 352(10): 987–996.

Tirosh, I., Venteicher, A. S., Hebert, C., Escalante, L. E., Patel, A. P., Yizhak, K., Fisher, J. M., Rodman, C., Mount, C., Filbin, M. G., Neftel, C., Desai, N., Nyman, J., Izar, B., Luo, C. C., Francis, J. M., Patel, A. A., Onozato, M. L., Riggi, N., Livak, K. J., Gennert, D., Satija, R., Nahed, B. V., Curry, W. T., Martuza, R. L., Mylvaganam, R., Iafrate, A. J., Frosch, M. P., Golub, T. R., Rivera, M. N., Getz, G., Rozenblatt-Rosen, O., Cahill, D. P., Monje, M., Bernstein, B. E., Louis, D. N., Regev, A. and Suva, M. L. (2016). “Single-cell RNA-seq supports a developmental hierarchy in human oligodendroglioma.” Nature 539(7628): 309–313.

Toms, S. A., Kim, C. Y., Nicholas, G. and Ram, Z. (2019). “Increased compliance with tumor treating fields therapy is prognostic for improved survival in the treatment of glioblastoma: a subgroup analysis of the EF-14 phase III trial.” Journal of neuro-oncology 141(2): 467–473.

Trepant, A. L., Bouchart, C., Rorive, S., Sauvage, S., Decaestecker, C., Demetter, P. and Salmon, I. (2015). “Identification of OLIG2 as the most specific glioblastoma stem cell marker starting from comparative analysis of data from similar DNA chip microarray platforms.” Tumour Biol 36(3): 1943–1953.

Voronkova, M. A., Luanpitpong, S., Rojanasakul, L. W., Castranova, V., Dinu, C. Z., Riedel, H. and Rojanasakul, Y. (2017). “SOX9 Regulates Cancer Stem-Like Properties and Metastatic Potential of Single-Walled Carbon Nanotube-Exposed Cells.” Scientific reports 7(1): 11653.

Voss, V., Senft, C., Lang, V., Ronellenfitsch, M. W., Steinbach, J. P., Seifert, V. and Kogel, D. (2010). “The pan-Bcl-2 inhibitor (-)-gossypol triggers autophagic cell death in malignant glioma.” Molecular cancer research : MCR 8(7): 1002–1016.

Wang, J., Xu, S. L., Duan, J. J., Yi, L., Guo, Y. F., Shi, Y., Li, L., Yang, Z. Y., Liao, X. M., Cai, J., Zhang, Y. Q., Xiao, H. L., Yin, L., Wu, H., Zhang, J. N., Lv, S. Q., Yang, Q. K., Yang, X. J., Jiang, T., Zhang, X., Bian, X. W. and Yu, S. C. (2019). “Invasion of white matter tracts by glioma stem cells is regulated by a NOTCH1-SOX2 positive-feedback loop.” Nat Neurosci 22(1): 91–105.

Wiranowska, M., Ladd, S., Smith, S. R. and Gottschall, P. E. (2006). “CD44 adhesion molecule and neuro-glial proteoglycan NG2 as invasive markers of glioma.” Brain Cell Biol 35(2-3): 159–172.

Yoshimura, Y., Shiino, A., Muraki, K., Fukami, T., Yamada, S., Satow, T., Fukuda, M., Saiki, M., Hojo, M., Miyamoto, S., Onishi, N., Saya, H., Inubushi, T., Nozaki, K. and Tanigaki, K. (2015). “Arsenic trioxide sensitizes glioblastoma to a myc inhibitor.” PloS one 10(6): e0128288.

